# Design principles of 3D epigenetic memory systems

**DOI:** 10.1101/2022.09.24.509332

**Authors:** Jeremy A. Owen, Dino Osmanović, Leonid A. Mirny

**Affiliations:** Department of Physics, Massachusetts Institute of Technology, Cambridge, MA, USA; Department of Mechanical and Aeronautical Engineering, UCLA, Los Angeles, CA, USA

## Abstract

The epigenetic state of a cell is associated with patterns of chemical modifications of histones (“marks”) across the genome, with different marks typical of active (euchromatic) and inactive (heterochromatic) genomic regions. These mark patterns can be stable over many cell generations—a form of epigenetic memory—despite their constant erosion due to replication and other processes. Enzymes that place histone marks are often stimulated by the same marks, as if “spreading” marks between neighboring histones. But this positive feedback may not be sufficient for stable memory, raising the question of what is. In this work, we show how 3D genome organization—in particular, the compartmental segregation of euchromatin and heterochromatin— could serve to stabilize an epigenetic memory, as long as (1) there is a large density difference between the compartments, (2) the modifying enzymes can spread marks in 3D, and (3) the enzymes are limited in abundance relative to their histone substrates. We introduce a biophysical model stylizing chromatin and its dynamics through the cell cycle, in which enzymes spread self-attracting marks on a polymer. We find that marks localize sharply and stably to the denser compartment, but over several cell generations, the model generically exhibits uncontrolled spread or global loss of marks. Strikingly, imposing limitation of the modifying enzymes—a plausible but oft-neglected element—totally changes this picture, yielding an epigenetic memory system, stable for hundreds of cell generations. Our model predicts a rich phenomenology to compare to experiments, and reveals basic design principles of putative epigenetic memory systems relying on compartmentalized 3D genome structure for their function.

## I. INTRODUCTION

Remembering gene expression states—that is, which genes are “on” or “off”—is a striking capability of living cells. In an appropriately organized biochemical system, the memory of a collective state can be dramatically longer-lived than any constituent molecule or interaction. For example, growing cells can stably remember an expression state encoded in the abundances of diffusible transcription factors (TFs) which regulate their own synthesis, even as any particular TF molecule is diluted away by cell growth. Such “epigenetic” memory systems been identified in nature [1], recapitulated synthetically [2, 3], and have also been subject to extensive theoretical study, which has begun to reveal the design principles [4, 5] for this kind of memory—for example the often-central role of ultrasensitivity [6] in enabling it.

But the chromatin of eukaryotes may play host to a qualitatively different sort of epigenetic memory, one held locally to genes in the physicochemical state of DNA or the associated histone proteins. Differential silencing of two identical copies of a gene, as seen, for example, in X-chromosome inactivation or the FLC locus of *Arabidopsis* [7], is a “smoking gun” [8] of such so-called *“cis*-memory” [9], because freely-diffusing molecules such as TFs would be expected to act on both copies in the same way. Chromatin-based mechanisms are thought to contribute broadly to the stability of differentiated cell fates in multicellular eukaryotes [10, 11].

Histones are subject to scores of covalent, post-translational modifications (“marks”), which vary across the genome in patterns correlated with gene expression. It has been suggested that these patterns could be the seat of a chromatin-based cis-memory of gene expression states [12]. However, chromatin is subject to huge physical and chemical disruptions through the cell cycle, raising the question of how such patterns are maintained. What kind of molecular order is needed—that is, how would the systems that add and remove histone marks have to work—to make memories out of mark patterns?

In answering this question, it seems likely we will need to go beyond the conceptual tools developed for memories based on diffusible factors. In particular, the biochemistry of histone modification, together with the fact that histones are embedded in chromatin, opens the door for the involvement of 3D genome organization in memory.

We focus on the best candidate for a chromatin-based memory system, which is found in the transcriptionally silent parts of the genome—the heterochromatin [13, 14]. Heterochromatin is rich in “repressive” histone marks—the lysine trimethylations H3K9me3 or H3K27me3—made by enzymes that are stimulated by their own products [15, 16]. This so-called “reader-writer” positive feedback can be thought of as “spreading” marks between nearby histones. Additionally, in heterochromatin, marked histone H3/H4 tetramers can be retained locally when the replication fork passes, partitioning stochastically between the old and new strands [17, 18] though (by necessity) they are diluted in the process by newly synthesized, unmarked histones. The combination of these two features is highly suggestive of a stable memory system, in which local mark spreading accurately restores mark patterns after their partial erasure at replication. However, simple mathematical models [19, 20] of this mechanism reveal a basic instability— if mark spreading is strong enough to restore a partially erased pattern, it will also spread ectopically, unless somehow constrained. Alternatively, a mark pattern may be sustained by external reinforcement, for example by “nucleation sites” or “genomic bookmarks” [19–21], but a pattern determined by such external influences is not itself a seat of memory.

Recent experiments suggest that reader-writer enzymes may be able to spread histone marks “in 3D” [16, 22, 23], that is, between histones that are nearby in space because of how chromatin is folded, not just “in 1D” along the chromatin polymer. A mechanism involving 3D spread could be sensitive to the 3D structure of the genome. Heterochromatin is “sticky”, exhibiting compartmentalization from, and a higher volume density than, other parts of the genome [24–29]. This raises the tantalizing possibility of a *bidirectional* coupling between the folding and the chemical state of the chromatin polymer [30–33]— patterns of histone marks might influence 3D genome organization, which might in turn influence the patterns of marks.

In this work, our goal is to understand whether a memory of mark patterns can be stably maintained by such a coupling *alone*. To do this, we introduce and study a simple model (Figure 1) of “sticky” self-attracting marks of single type spreading on a polymer. Several studies have explored the consequences of this kind of bidirectional coupling using similar biophysical models [33–36]. However, broadly, these works had difficulties maintaining stable *patterns* of marks, exhibiting a clear tendency towards only a bistable memory of a globally homogeneous marking state (i.e. high marking or low marking), as seen also in the seminal model of Dodd et al. [37], which had no feedback of the mark pattern onto the 3D structure. Our goal, by contrast, is to find a mechanism where bidirectional coupling between marks and 3D structure enables stable, self-sustaining memory of a mark pattern.

**FIG. 1.**
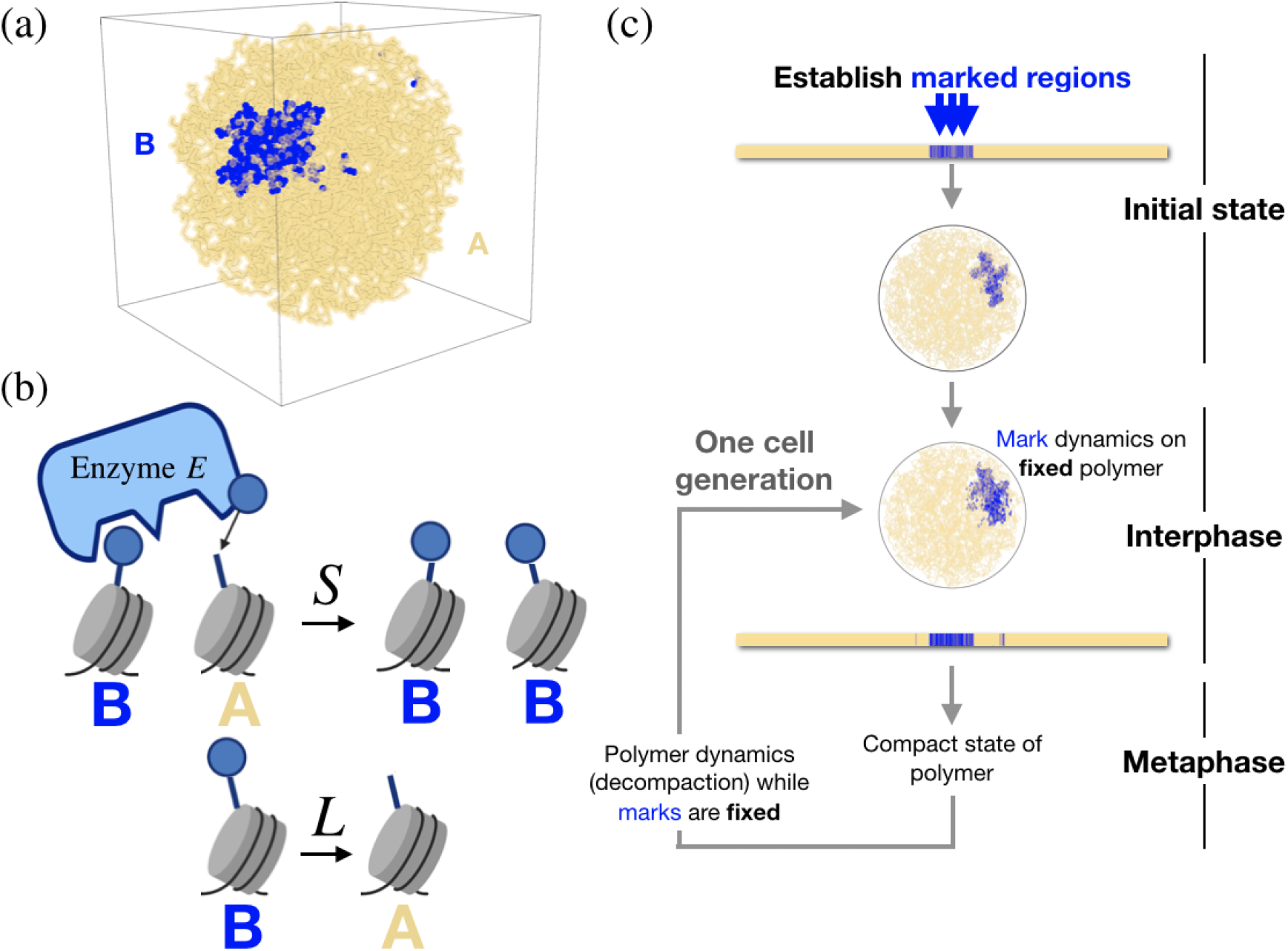
Our model. (a) Chromatin in the nucleus modeled as a spherically confined copolymer with monomers of two types, A (pale yellow) and B (blue), representing a varying pattern of histone marks. Monomers of type B, which represent regions bearing heterochromatic marks, self-attract. (b) Marks spread to neighbors (in 3D space, within an interaction radius), at rate *S*, and are lost everywhere uniformly at rate *L*. (c) The overall dynamics of our model consist of alternating phases of polymer dynamics and mark dynamics, coarsely stylizing the cell cycle.

Using our model, we show that just like models of 1D spreading [19], 3D spreading of self-attracting marks appears to possess a fundamental instability—an all-or-none tendency to either global loss or uncontrolled spread of marks. We find that the instability is a consequence of the questionable assumption that the enzyme responsible for spreading marks is in excess relative to its histone substrates. We then show that limitation of the spreading enzyme—very plausible biologically [38, 39]— stabilizes mark patterns, in this 3D epigenetic memory system, for hundreds of cell generations, and yields a rich phenomenology to compare to experiment.

The memory effect we uncover depends critically on three key ingredients:

1. **strong self-attraction** of marked regions, leading to compartmentalization and a two to threefold densification of marked regions,
2. **3D spread** of marks, and
3. **limitation of the “reader-writer” enzyme** responsible for spread, relative to its histone substrates.

Together, these elements amount to basic design principles for epigenetic memory systems that exploit 3D genome structure for their function.

## II. MODEL

We developed a simple model coupling the 3D structure of chromatin—and its reorganization through the cell cycle—to the dynamics of a single epigenetic mark.

We model chromatin as a polymer, of *N* = 10^4^ monomers, confined within a sphere (Figure 1a). This may be viewed as representing a chromosomal region (e.g. a 2 Mb region, if each monomer represents a nucleosome) or, for example, a single “territorial” chromosome (if each monomer represents a 10 kb region).

The monomers in the polymer can be in one of two states—A or B—with B monomers representing marked, heterochromatic regions and A monomers representing (unmarked) euchromatic ones. All monomers experience short-ranged repulsion (excluded volume), but to reflect the “stickiness” of heterochromatin, B monomers experience a short-range attraction to other B monomers of magnitude *α* (in units of *k_B_T*) within an interaction radius *r_c_*, which we choose to be 1.5 times the monomer diameter.

To model the spreading of marks (Figure 1b), we suppose that A monomers turn into B monomers at a rate *S_n_B__*, where *n_B_* is the number of neighboring B monomers within the interaction radius *r_c_*, and *S* is a constant— the spreading rate. Importantly, we suppose throughout that spreading of marks can happen between monomers that are physically proximate in space, not just between nearest neighbors along the polymer chain.

To model the loss of marks, we suppose that B monomers turn back into A monomers at a constant rate *L*, uniformly at all sites. We intend this to coarsely represent processes that remove histone marks, including their loss by dilution with newly synthesized, unmarked histones when DNA is replicated. However, since replicational dilution occurs once through the cell cycle (occuring in *S* phase), it is not at all clear it can be modeled, even roughly, as loss taking place at some constant rate. To address this we also consider a variant of our model where loss happens all at once, periodically, instead of at a constant rate (Figure 4b and Figure S3). Our core results will prove insensitive to this detail.

We note that the spreading and loss dynamics we posit are identical to those of an Susceptible-Infected-Susceptible (SIS) epidemic model on a contact network [40]. The monomers of our polymer are like individuals whose “social” contact network is defined by the polymer configuration (and the interaction radius *r_c_*). The marked state is equivalent to the infected state in the SIS model. Later, we will use this analogy to better understand how density differences in the nucleus could contribute to epigenetic memory.

The final ingredient in our model is an *extremely stylized* representation of the cell cycle (Figure 1c). We suppose it consists of two alternating phases—an “interphase” during which the polymer is fixed in a single configuration while marks are spread and are lost, reaching a steady-state, and a “metaphase” during which the polymer is compacted into a condensed state and then refolds into a new “interphase” state, all while the marks are fixed. Repeated alternation of these phases constitutes the cell cycle dynamics of our model. A single round of polymer relaxation followed by mark dynamics we will refer to as one “cell generation”. that pattern. We then freeze it in that 3D configuration. What is the behavior of the mark dynamics on the resulting frozen polymer?

In the following sections, we will investigate the behavior of this model, and in particular its capacity under various conditions to exhibit a long-lived memory of mark patterns. What do we mean by “memory”? When we run the dynamics of our model, we must begin with; an *initial pattern* specifying the identities (A or B) of all the monomers. This initial pattern of marks then evolves over one or many cell generations. For some time, this evolving pattern will tend to resemble the initial one—this is memory.

## III. RESULTS

### A. Marks localize to dense regions providing stable, memory for one cell generation

We start by considering memory of the mark pattern over a single generation. We set an initial A/B pattern of marks, allowing the polymer to fold (that is, to relax in the potential set by the self-attraction of the marked, B, regions) forming a compartmentalized state set by that pattern. We then freeze it in that 3D configuration. What is the behavior of the mark dynamics on the resulting frozen polymer?

We see a striking phenomenon, which is that the mark dynamics approaches a steady-state pattern which closely resembles the initial mark pattern used to fold the polymer (Figure 2a and 2b). As long as the spatial polymer configuration remains unchanged, this steady state pattern is stable to huge perturbations, such as complete randomization of the pattern (every monomer randomly set to the A or B state, Figure 2a) or wholesale erasure of half of the pattern (Figure 2b). The reason for this stability is that the marks, which spread in 3D, tend to localize to the spatially dense compartment that was formed by the originally marked regions, due to their self-attraction. It is as if the mark pattern has been “memorized” in the 3D configuration of the polymer.

**FIG. 2.**
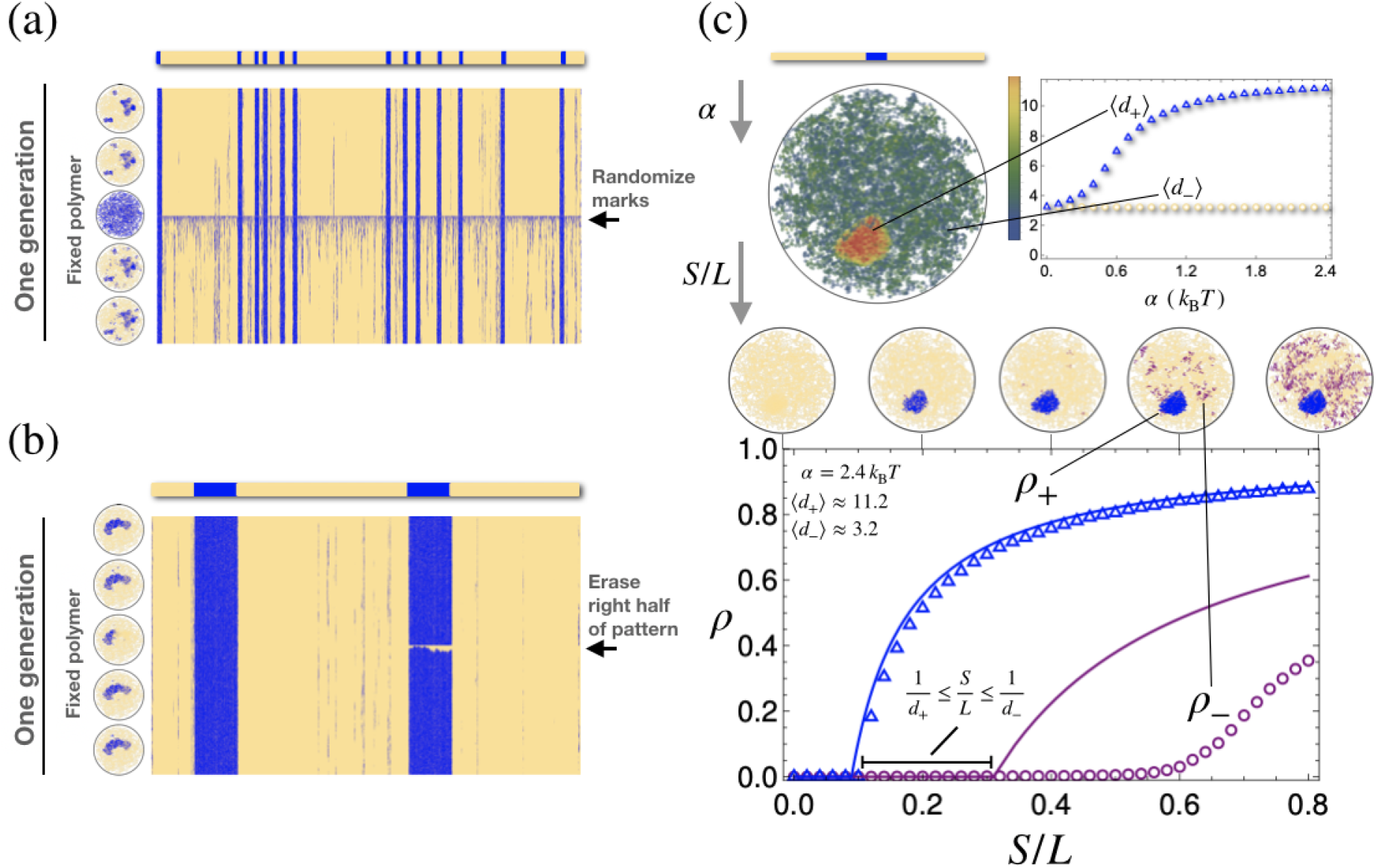
Mark dynamics and density differences. (a) The mark dynamics over time (with *S/L* = 0.5) on a fixed polymer initially folded according to an initial pattern (with *α* = 2.4 *k_B_T*). Time advances from top to bottom. The steady state closely resembles the initial pattern and is stable to huge perturbations (e.g. randomization of the mark pattern). Inset circles (left) show snapshots of the state over time. The polymer is frozen throughout. (b) The same behavior as shown in panel (a) with a different initial pattern and a different kind of perturbation. (c) Top: when the polymer folds, marked regions tend to be denser (red) than unmarked ones (green), due to the self-attraction of marks. Bottom: in turn, when marks evolve according to their dynamics of spreading and loss, they tend to localize in dense regions. In the steady state, the linear mark density *p* (purple circles and blue triangles) depends on *S/L*. In dense regions (blue), the relationship is given very closely by a simple mean-field theory (solid lines).

To understand this effect systematically, it is helpful to consider the fate of a single, contiguous domain of B monomers (“marks”) against a background of A monomers (Figure 2c). To quantify the steady-state localization of marks, we will compare the steady-state marking fraction *ρ*_+_ (fraction of monomers that are marked) inside the initially marked domain to the steady-state marking fraction *ρ*_-_ outside the domain. Perfect recovery of the initial mark pattern would correspond to *ρ*_+_ = 1 and *ρ*_-_ = 0. This never happens in our model, but simulations reveal that for a given choice of *α* (and so the density difference), there is a broad range of values of *S/L* for which *ρ*_+_ > 0 and *ρ*_-_ ≈ 0 (Figure 2c, bottom). This corresponds to sharp localization of marks to the dense compartment.

An analogy to epidemic spreading helps us understand how this localization phenomenon depends on the difference in the spatial density of the compartments. The density difference—which depends on the strength of self-attraction *α*—can be quantified by comparing the average number 〈*d*_+_〉 of neighbors of a monomer in the dense region, with that of a monomer in the diffuse region, 〈*d*_-_〉. The mark spreading dynamics of our model are identical to a Susceptible-Infected-Susceptible (SIS) epidemic model on a network [40]. A simple mean field approximation (Appendix A) for this model relates the steady-state probability of a monomer being marked *ρ* to the number d of its neighbors: *ρ* ≈ max(1 – *L*/(*Sd*), 0). The validity conditions for this approximation suggest that at least when *S/L* lies in the range:

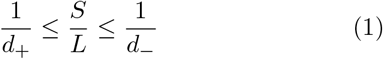

there will be sharp localization of marks to the dense region, with very few marks in the diffuse region (*ρ*_+_ > 0 and *ρ*_-_ ≈ 0). This range grows broader as the difference in the density of the compartments grows. Also, the marking fraction *ρ*_+_ at the upper end of this interval (when *S/L* = 1/*d*_-_) is given in the mean field approximation to be 1 – *d*_-_/*d*_+_, which approaches the case of perfect recovery (*ρ*_+_ = 1) as the ratio of the densities grows. These facts highlight how the difference in density between heterochromatin and euchromatin could play an essential role in the localization of spreading marks to the heterochromatin.

With increasing α, the monomers in the dense region become tightly packed and the densities saturate around 〈*d*_+_〉 = 11.2, 〈*d*_-_〉 = 3.2, yielding a density ratio of around 〈*d*_+_〉/〈*d*_-_〉 ≈ 3.5. Simulations (Figure 2c, bottom) of this case show that localization of marks occurs in a range of *S/L* that includes the interval (1), as predicted, providing robust recovery of the initial mark pattern, within one cell generation.

Although the large-scale pattern of the marks is preserved over one generation, there is never perfect recovery of the initially fully marked domain, because the steady-state marking fraction *ρ*_+_ is always less than 1. What happens, then, if we iterate? If we refold the polymer, as the marking fraction is lower, there are fewer self-attracting monomers in the domain, and the spatial density will drop in the next iteration. However, if *S/L* is large enough the marking fraction in the marked domain can converge to a positive marking fraction 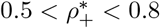.

Taken together, these results show that a density difference between the compartments provides sharp localization of marks to the dense phase. The pattern of marks established through this localization is stable to erasure and noise in marks, and does not require fine-tuning of the spreading rate. Additionally, if spreading is strong enough, the mark density in a dense region can sustain itself even if the polymer refolds over multiple generations. However, we did not consider what happens *globally* to the pattern of marks along the polymer over multiple generations.

### B. Memory is lost over multiple cell generations

We will now study the evolution of the whole mark pattern over multiple generations, starting again from an initial pattern consisting of a small contiguous marked domain against a background of unmarked monomers. An example is shown in Figure 3a—time (measured in generations) advances from the top to the bottom, and each row represents the mark pattern at the corresponding point in time. In this example, within tens of generations, marks spread across the whole polymer.

**FIG. 3.**
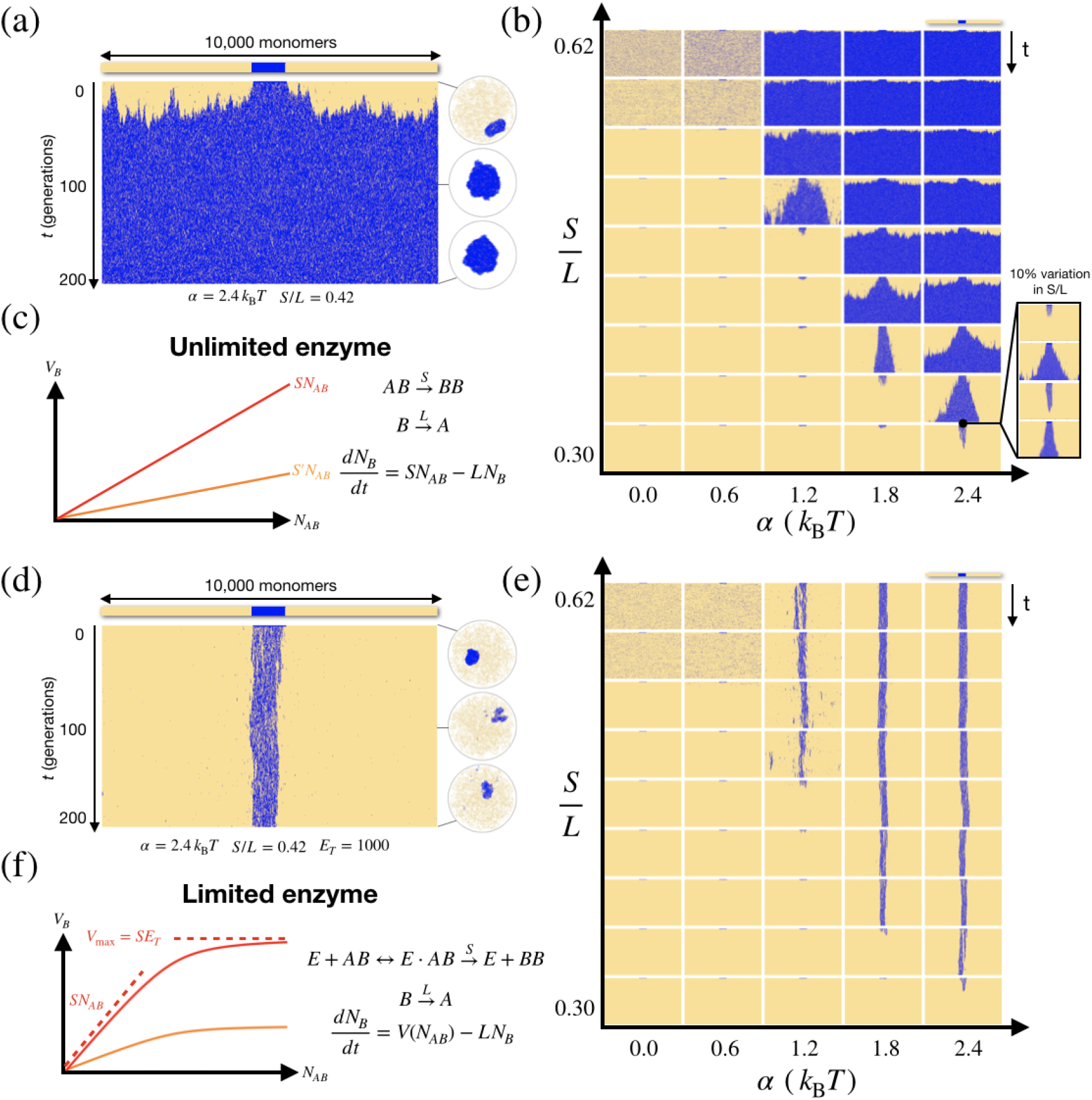
Memory over multiple generations. (a) Time evolution of a mark pattern over 200 generations, starting from an initial pattern consisting of a single domain of 1000 marked monomers. Inset circles show snapshots in time of the polymer configuration. For this choice of parameter (*a* = 2.4 *k_B_T, *S/L** = 0.42) marks spread everywhere and the polymer collapses. (b) Time evolution of mark pattern as a function of *α* and *S/L*. (c) In our model without limited enzyme, the global marking rate in the nucleus *V_B_* is proportional to the number *N_AB_* of A-B pairs. (d) As in (a), but with limited enzyme, *E_T_* = 1000. (e) As in (b), but with limited enzyme, *E_T_* = 1000. (f) In our model with limited enzyme, the global marking rate in the nucleus *V*_B_ is proportional to the number *N_AB_* of A-B pairs when it is small, but then saturates at a value of *V*_max_ = *SE_T_*.

Sweeping through the parameter space of our model (Figure 3b), what we find is an unstable, all-or-none behavior. When *S/L* is greater than a critical value *λ_c_*(*α*) (which depends on *α*), the marking fraction in the originally marked domain quickly stabilizes, but the marked region grows in spatial extent, spreading until it covers the whole polymer. When *S/L* is less than the critical value, the marks are instead lost globally. In both cases, memory of the initial state is lost within tens of generations.

Only when *S/L* is fine-tuned—set very close to the critical value—can there be a long-lived memory of the initial state. With fine-tuning, it is possible to achieve memory that lasts hundreds of generations, although even in this case, there is a plain tendency towards uncontrolled spread or global loss over that timescale. The same basic instability is apparent in the closely related model of Sandholtz et al. [36], who found that fine-tuning of parameters was required to achieve even just 5 generations of mark pattern memory.

### C. Enzyme limitation stabilizes epigenetic memory

The uncontrolled spreading of marks we describe above depends fundamentally on a feedback in which, as there are more and more marks, there is also a higher and higher global modification rate *V_B_* (the total rate at which marks are created in the system). In the model as we have described it so far, this is easy to envision, because *V_B_* is simply proportional to the total number *N_AB_* of A-B monomer pairs (within the interaction radius of one another), across the entire polymer (Figure 3c). It is natural for this to increase with the total number of marks *N_B_*, enabling uncontrolled spreading of marks.

However, histone marks do not spread themselves— spreading requires the action of a reader-writer enzyme. If this enzyme is limited in abundance, it effectively sets the maximum global modification rate in the nucleus— intuitively, this is the *V*_max_ of the enzyme. We will see that this creates a kind of negative feedback that can dramatically stabilize the memory of the initial mark pattern (Figure 3d, e).

To account for the limitation of the reader-writer enzymes we must introduce a model of their enzymology. Perhaps the simplest choice is a Michaelis-Menten-type scheme [41, 42] where A-B pairs that are within the interaction radius act as the substrate (Figure 3f):

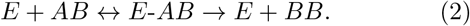

Under a few additional assumptions (Appendix B), adding this to our model yields a new model of the just the same form we had above (before introducing the enzyme), but with an effective spreading rate *S*_eff_, given by:

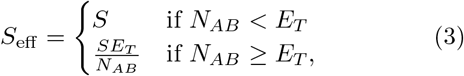

where *N_AB_* is the total number of A-B spatial pairs (within the interaction radius of one another), and *E_T_* is the (constant) total amount of enzyme (bound and unbound). Note that this means that when the quantity of substrate (*A-B* pairs within the interaction radius) increases sufficiently so as to saturate the enzyme (as it does when *S/L* > *λ_c_*(*α*)), the global marking rate will max out at “*V*_max_” = *SE_T_*. This means that when *S/L* exceeds the critical value, the steady-state average number of marks 〈*N_B_*〉 will be *SE_T_/L*, which plainly rules out the uncontrolled spreading of marks seen in the unlimited enzyme case (Figure 3a). And critically, the position of an initially imposed marked domain is remembered (Figure 3d, Figure S2) over many generations. This finding is seen across a broad range of parameters, as long as self-attraction is strong enough, and does not require fine tuning (Figure 3e). Stability of the mark pattern is also seen when loss occurs purely by replicational dilution (modeled as random loss of half the marks) once every cell cycle period *T*_div_ instead of at a constant rate *L*. The behavior of this “dilution-only” model appears similar qualitatively, and can be matched quite closely to a constant loss model with *L*_eff_ = log(2)/*T*_div_ (Figure 4b, Figure S3).

**FIG. 4.**
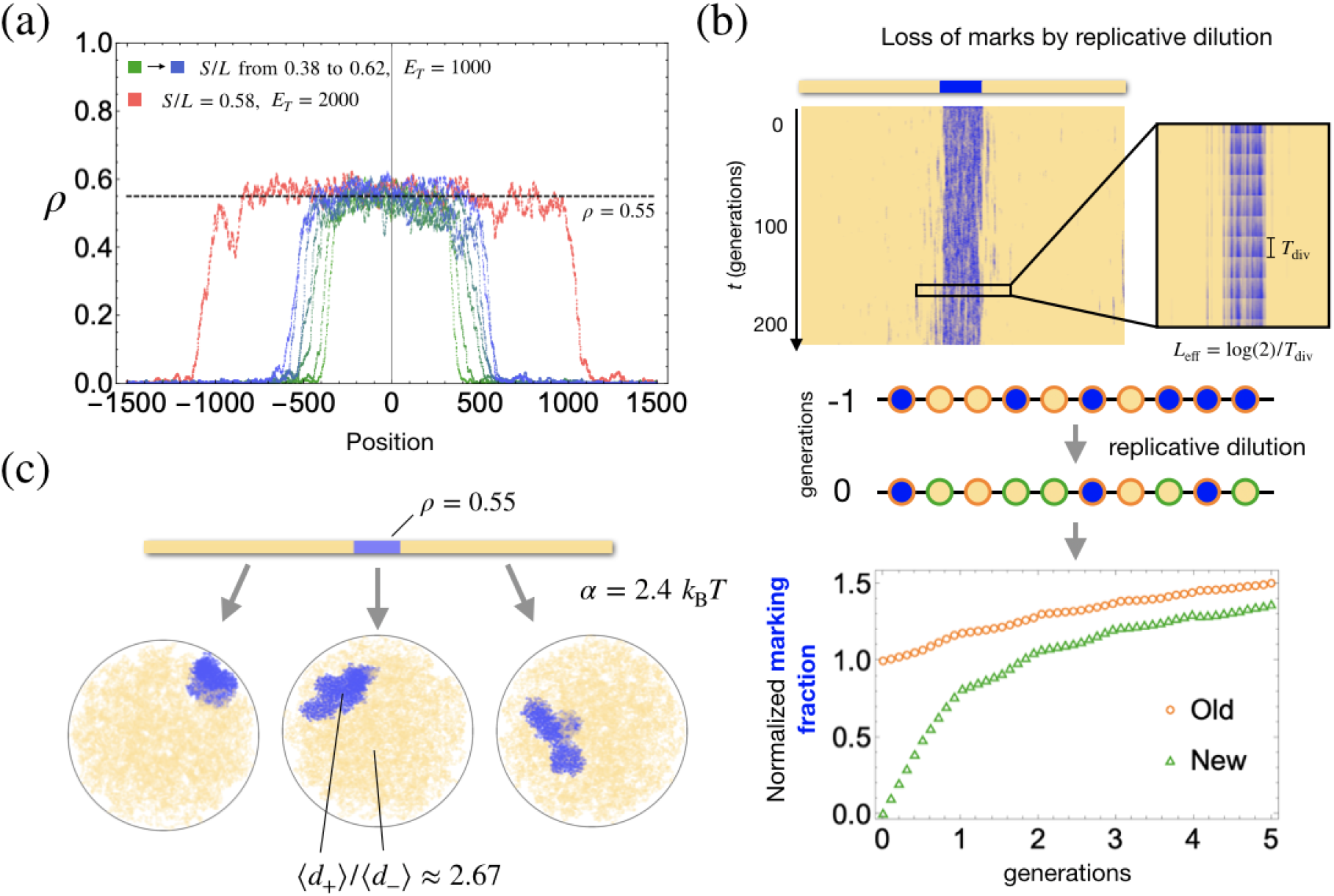
The character of semimarked domains. (a) Centered profiles of stable domains arising in our model, averaged over many generations, showing that they are consistently *semimarked*—with a stable marking fraction of about *ρ* = 0.55. This marking fraction is insensitive to change in *S/L* or *E_T_*. (b) In a variant of our model where loss is purely by replicative dilution, we keep track of the marking fraction of the “new” monomers (green) that were replaced by dilution at a given time, losing their marks, and compare it to the marking fraction for the monomers that were not replaced (orange). The marking fraction (normalized to its value at the initial time) grows for both populations, a signature of semimarking. Compare, e.g. to Figure 3, panel (e) of [44]. Note when tracked over several generations, the old and new monomers accrue marks even though the average marking fraction across the domain is constant, because these two populations become an ever smaller fraction of the whole. (c) Examples of the 3D shape of semimarked domains, illustrating their tendency to be aspherical.

This system can be viewed as “self-tuning” to a critical value. If *S/L* > *λ_c_*(*α*) but there is limited enzyme, the average number of marks neither grows indefinitely nor shrinks to zero, but rather approaches a positive, stable value. In the original, “unlimited enzyme” model this cannot happen except (possibly) if *S/L* = *λ_c_*(*α*). Intuitively, this suggests it may be fruitful to think of the enzyme as actively tuning *S*_eff_/*L* to the critical value *λ_c_*(*α*). Taking this idea seriously, we should therefore expect that

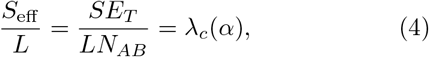

while *N_B_* = *SE_T_/L*, so that *N_AB_/N_B_*, which is a sort of surface area to volume ratio, should equal 1/*λ_c_*(*α*). Our simulations suggest this intuitive relation is satisfied very closely (Figure S1). We note that the same “tuning to criticality” has been observed in the conserved contact process [43], which is a 1D model closely related to the mark dynamics of our model in the limited enzyme regime.

Together, these findings suggest that to maintain memory over many cell generations, the enzyme should be limited relative to its substrate.

### D. Stable domains are semimarked

The stable mark domains (Figure 3e) that emerge in the limited enzyme regime have a striking property—as *S/L* or *E_T_* is varied, the total number of marks changes, but the *marking fraction* within the domain remains roughly constant, around *ρ* = 0.55 (Figure 4a). To accommodate different numbers of marks, domains vary instead in their linear extent.

The “semimarked” character of the domains that emerge in the limited enzyme regime has a number of consequences that could be experimentally observable.

First, Alabert et al. [44] found experimentally that certain histone PTMs require several cell generations to be fully established on new histones after replicational dilution. This slow establishment is invariably accompanied by increasing modification levels on “old” histones, and it is therefore a signature of the semimarking predicted by our model.

To emulate these experiments, we used the version of our model in which loss occurs purely by replicational dilution and tracked the recovery of marks after dilution (Figure 4b). As Alabert et al. saw in their experiments, we also observe “slow, perpetual” marking of newly deposited histones (i.e. monomers that lost their marks at a given time) over many cell generations, along with continued marking of the old histones (monomers that did not lose their marks). Continuous gain of marks on the old histones, in both simulation and experiment, indicates that some were unmarked to start off—a reflection of the semimarking that emerges naturally in our model.

Second, semimarking may have implications for the 3D structure of marked domains. Semimarked domains fold into structures that are less spatially dense than they would be if they were fully marked, leading to a density difference of about 2.67-fold, which is close to what has been reported for the difference between heterochromatic and euchromatic regions in the nucleus [28]. Additionally, in our simulations, when marked domains fold they tend not to be spherical, and they are not homogeneous but exhibit substructure, with a dense interior and a surface consisting of unmarked, intercalary stretches of monomers.

### E. Marks redistribute over time

Figure 3e shows that in the limited enzyme version of our model, in the regime of strong self-attraction, mark patterns consisting of a single contiguous domain can be remembered, at least for hundreds of generations. We want to emphasize that the memory reflected in Figure 3e *is of the overall position of the domain*, rather than of the marking of any individual monomer, or of the size of the domain (which is set by *E_T_*). In Figure S2 we illustrate this positional memory explicitly by showing the fate of domains with different initial positions along the polymer. The eventual loss of this positional memory is by extremely slow diffusion of the position of single domains.

If the initial pattern is more complex, consisting of multiple, noncontiguous marked domains, it can be remembered for a time that depends on the pattern. When domains are large, equally-sized, and well-separated, they can be remembered for hundreds of generations (Figure 5a). However, over a longer timescale, separate domains compete with one another for the shared enzyme pool, and also merge when they are close along the polymer. The final outcome of this process is always a single contiguous domain. We term this phenomenon “mark redistribution”, since it involves a change in the pattern of marks along the polymer while the total (average) number of marks remains fixed.

**FIG. 5.**
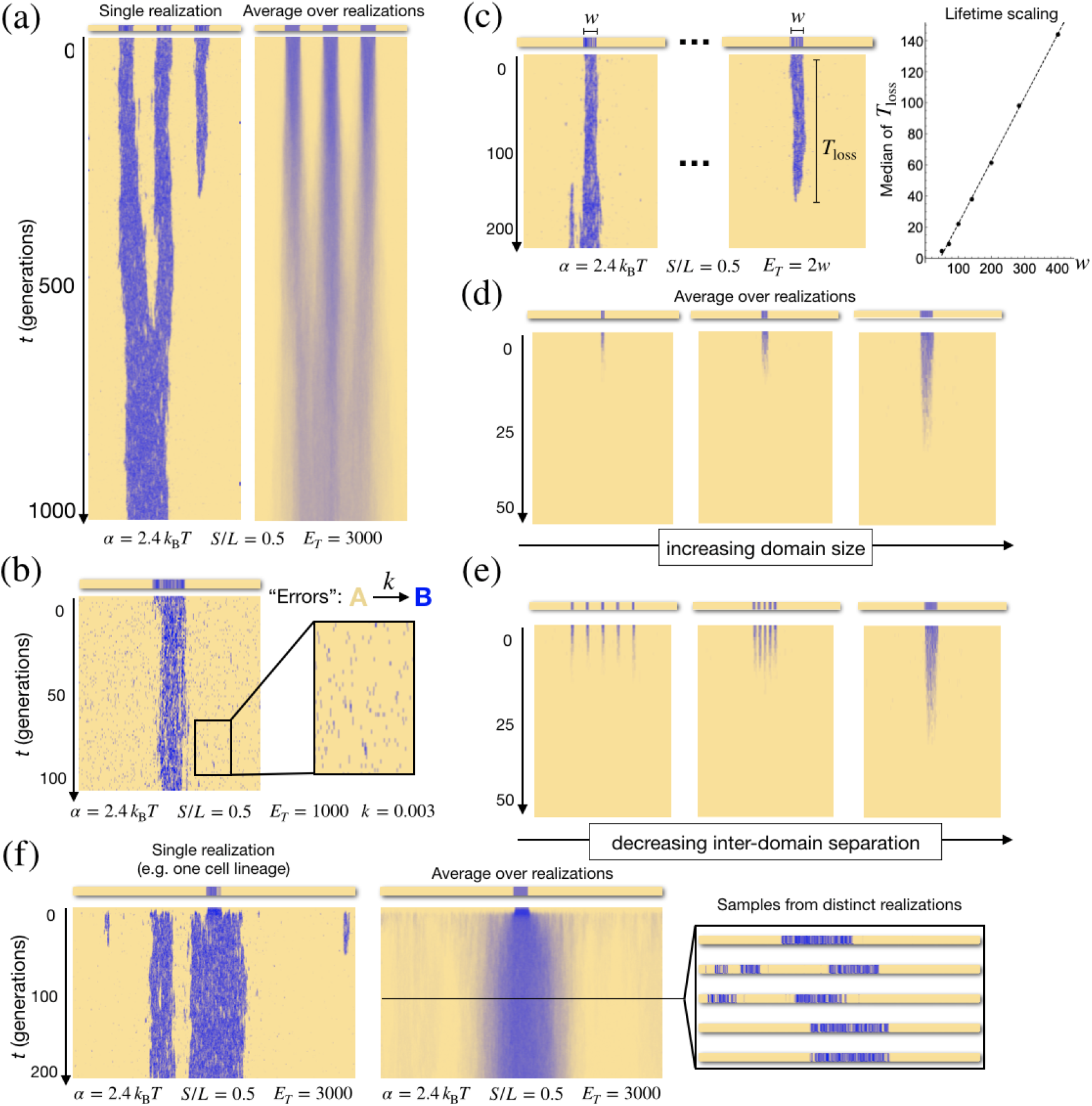
Dynamics of mark redistribution in the limited enzyme regime. (a) Time evolution of a pattern consisting of three, equally-sized mark domains. The pattern is stable for hundreds of generations, but in any given realization (left), marks eventually redistribute to form a single contiguous domain. In the average of many realizations (right) the three domains appear instead to merge. (b) Tiny “domains” introduced by spontaneous marking at a rate k are lost immediately—a form of error correction. (c) Two equally-sized domains a long way apart compete for limited enzyme pool. (d) A small domain competing with a much bigger one (not pictured) lasts longer, the bigger it is. (e) Multiple small domains competing with a much bigger one (not pictured) last longer the closer they are together. (f) Expansion of marks from a small initial domain discloses striking differences between single realizations and the average over many.

Mark redistribution could be seen as purely detrimental to the preservation of a particular mark pattern, but it can also be viewed as a form of error correction. When “errors” are continuously introduced in the form of monomers spontaneously switching from A to B— creating very small “domains” consisting of individual monomers—these errors are destroyed immediately by redistribution of the marks to a larger domain (Figure 5b). In other words, in the presence of spontaneous switching, the same mechanism which gives rise to the gradual loss of memory over long timescales is necessary for its preservation over shorter timescales. Mark redistribution can provide stability to a pattern against random errors.

To explore the phenomenon of mark redistribution more quantitatively, we ran simulations of equally-sized domains kept totally isolated from one another (never coming into 3D contact), but which share the same enzyme pool. These widely separated domains compete for the limited enzyme. Invariably, one of the domains is eventually lost, with the median time until loss increasing with domain length *w*. The lifetime appears to grow linearly with w, above a minimum length of roughly 50 monomers (Figure 5c). This finding deviates from the quadratic scaling we would expect if the size of the domains were undergoing unbiased diffusion—once there is any significant difference in the size of the two domains, there is a strong tendency for the larger domain to grow at the expense of small ones.

Similar to the case of equally-sized domains, when a small domain competes with a much larger domain, we see that the larger the small domain is, the longer it lasts (Figure 5d). We also investigate the fate of multiple small domains (“islands”), also competing with a much larger domain (not pictured). We find that the smaller the distance between the islands, the longer they last before being lost at the expense of the growth of the much larger domain (Figure 5e).

Finally, in Figure 5f, we consider the case in which marks are initially present in a small region and then *S/L* or *E_T_* is suddenly increased (e.g. in the course of development, or as an experimental perturbation, see Discussion). This expansion process reveals striking differences between population averages and single realizations of our model. In the population average (i.e. the average over many stochastic realizations), the initially marked domain appears to expand linearly into a larger one. However, looking at single realizations (individual “cell lineages”) reveals that in fact, new marks emerge in non-contiguous stretches randomly located near the initial marked domain. The resulting pattern is then remembered for some time in individual lineages. Over hundreds of generations, in each lineage, marks redistribute to form one or a few large domains, strongly biased to include the small initially marked domain— explaining the population averaged picture of linear expansion of the original domain. With single-cell epigenomics gaining momentum [45], such predictions regarding cellular heterogeneity in mark patterns may soon be experimentally testable.

## IV. DISCUSSION

The key challenge for any account of how cells remember is to explain how the memory is preserved when cells grow and divide. Genetic information is preserved due to the self-templating nature of DNA and the intricate choreographing of DNA replication and cell division. For a putative epigenetic memory held in histone marks, the “spreading” of marks mediated by “reader-writer” enzymes has emerged as a central motif. This mechanism is highly evocative of a stable memory, since spreading could restore mark patterns partially effaced by replicational dilution. However, mathematical models suggest that simple mark spreading, both in 1D models of linear spread [19], as well as in models incorporating 3D spread, is fundamentally unstable—if spreading is strong enough to sustain marks against loss, they will tend to spread uncontrollably. If marks are to serve as a memory of cellular state, something must serve to prevent uncontrolled spread of marks.

Prior authors have addressed this problem in a number of ways. Many models, such as the seminal, early work of Dodd et al. [37] or the recent work of Katava et al. [35], sidestep the issue by modeling only a small chromatin region. The “all-or-none” character of spreading we have described above features in these models, and so the memory in these models is *bistability—*they remember a single bit, not a mark pattern. Many such bistable memory elements arrayed across the genome could yield a capacious epigenetic memory, but only if they can be assumed to be independent of one another, e.g. implicitly isolated from one another by boundaries. Other models, such as [46], include boundaries more explicitly. The model of Erdel et al. [34] can also be interpreted as positing boundaries, with the “conditional nucleation sites” serving as the bistable elements, and the regular sites between them serving as boundaries.

In theory, 1D mark spreading with perfect boundaries separating domains could serve as an exceptional memory system, because in principle, memory would be lost only when marks spontaneously vanished in a domain, due to many mark loss events happening one after another by chance, before a spreading event intervenes. In the 1D model with large *S/L*, loss of memory by this mechanism is exponentially rare in the domain size ([47], p. 71). However, the memory is very fragile to any “leakage” of the boundaries creating ectopic marks. Leakage could occur through occasional 3D spread, porosity of the 1D boundaries, or “unrecruited” histone modification by writer enzymes.

If 3D spread is possible, the structure of the genome in space comes into play [31–33]. Several prior models explored the possibility that 3D structure could stabilize a mark pattern in the context of 3D spread, but in contrast to our work, these studies did not find a lasting memory of an initial mark pattern—exhibiting instead patterns determined by external influences (e.g. a static 3D contact structure in the case of [48]), or requiring fine-tuning of parameters to achieve even a short-lived memory, such as in the model of Sandholtz et al. [36], perhaps the closest to our own.

Here, we show that the solution to this fine-tuning problem is to add enzyme limitation to this system. The combination of: (1) self-attraction of marks leading to a formation of a denser, marked compartment, (2) 3D spread of marks, and (3) enzyme limitation, provides a very stable memory of the genomic position of a domain of marks. All three of these ingredients are required for this effect.

Intuitively, our mechanism relies on the encoding of memory in different forms in different phases of the cell cycle. In interphase, memory is held in the 3D structure of the genome, in the form of density differences, because marks sharply localize to dense regions. In metaphase, when the 3D structure is being totally reorganized, memory is held in the 1D sequence of marks. Enzyme limitation prevents the uncontrolled spread or global loss of marks, by fine-tuning the effective spreading rate to a critical value that keeps the total number of marks constant.

Limited enzyme is very plausible biologically. First, it is compatible with estimates of the physiological concentrations of relevant species. For example, the concentration of nucleosomes in the nucleus (hundreds of micromolar) is probably at least a hundred times that of H3K27 methyltransferase PRC2 (estimated to be less than 1 micromolar [49, 50]). Second, there is a long history of experiments suggesting the machinery assembling heterochromatin is limited (often referred to as a “titration” effect [39]). The position-effect variegation phenomenon in *Drosophila*, for example, where a gene translocated near heterochromatin gets silenced by the spreading of heterochromatin, is suppressed in the presence of extra copies of the (mostly heterochromatic) Y chromosomes [51]. A more recent example is the fact that the BAF complex both promotes and suppresses Polycomb repression, interpreted by Weber et al. [52] as reflecting genomic redistribution of limited Polycomb group proteins.

Our model makes a number of testable predictions. First and most basically, the model predicts that perturbations of 3D genome structure can impact the patterns of spreading marks. For example, reducing the spatial density of a chromatin region or relocating a locus to a more diffuse region should, by compromising 3D spread, reduce the marking fraction. The relationship between density and the marking fraction is nonlinear, and at a critical density (which depends on *S/L*), spreading cannot be sustained, and the marking fraction goes to zero (Figure 2c, Appendix A).

Second, our model predicts that changing the concentration E_T_ of (the active form of) an enzyme that spreads a certain mark will tend to have a linear effect on the (average) total number of those marks in the nucleus. Enzymological details could modify this relationship, but if the total number of marks were found to be insensitive to the concentration, it would suggest a system that was not enzyme limited.

Our model also predicts that if the ratio of spreading to loss (S/L) is below a critical value, the number of marks will catastrophically collapse, no matter how much enzyme is present. It is probably hard to control *S* experimentally since it is a microscopic, kinetic parameter, but for marks lost primarily by replicational dilution, the effective loss rate (per unit time) can be decreased by lengthening the cell cycle, providing an avenue to test this prediction.

Third, our model predicts semimarking—a marking fraction of around one-half—in stably marked regions, independent of the enzyme concentration E_T_. We believe a signature of this has already been seen in the experiments of Alabert et al. [44], with our model able to recapitulate some of their observations (Figure 4b). This prediction could be tested directly by any technique enabling absolute quantification of the prevalence of marks (fraction of histones bearing a mark) in heterochromatic regions.

Fourth, our model predicts that longer mark domains are more stable than shorter ones (Figure 5c, d). This could be tested by observing the fate of artificial ectopic mark domains of different lengths [34, 53, 54]. Very small domains that appear to exhibit a *cis*-memory of silencing [55] may rely on alternative mechanisms [56].

Finally, our model predicts that under certain conditions, such as the expansion of marks from a small initially marked region, significant cell-to-cell variability in mark patterns may develop, which may be heritable in individual cell lineages, but which is hidden in population averages (Figure 5f). This effect arises because the initially marked region grows differently in each cell, but is then remembered.

We believe our model reveals important design principles for epigenetic memory systems exploiting 3D genome organization. At the same time, the model has a number of important limitations, being a simplification as it is of a much more complicated biological reality.

For example, it may be too great a simplification to seek a chromatin-based memory involving a single spreading histone mark. In reality, there are many histone marks which can spread, compete directly (excluding one another), or interact in complex, indirect ways (for example, by influencing transcription [57], or replication timing [58]). And histone PTMs may also be coupled to other aspects of the chemistry of chromatin, such as patterns of DNA methylation or histone variants [59]. Interaction between multiple “epigenetic degrees of freedom” could enable behaviors that cannot be captured, even qualitatively, in a model involving a single histone mark.

Our model of the cell cycle is also extremely crude. In particular, we do not double the amount of chromatin during the cell cycle (e.g. as if to model *S* phase), and we suppose that chromatin does not move in interphase, which may be a tolerable idealization at large scales, but is certainly wrong at small scales [60, 61]. Whether these details affect our story qualitatively remains to be seen.

We want to conclude by emphasizing that any *cis*-memory mechanisms at work in eukaryotes seem to interact strongly with diffusible factors (such as transcription factors). This is crisply demonstrated, for example, by the experiments on nuclear transplantation and reprogramming. Transplantation of nuclei from differentiated tissues into enucleated oocytes can sometimes yield viable eggs, suggesting that memory of cell type can be totally erased by diffusible factors in the ooplasm [62, 63]. By investigating putative chromatin-based mechanisms as if they worked in isolation—we are not intending to suggest that they do work in isolation. What we think, rather, is that understanding the autonomous capacities of chromatin-based memory systems may prove invaluable to understanding the chemical systems cells use to remember, of which they are a part.

## ACKNOWLEDGMENTS

This work was supported by the NIH Center for 3D Structure and Physics of the Genome of the 4DN Consortium (U54DK107980) and NIH grant GM114190.

## Appendix A: Mean-field theory for mark dynamics on a spatial network with a dense region

Consider our mark dynamics—or completely equivalently, an SIS (Susceptible-Infected-Susceptible) epidemic model—on a (undirected) graph *G* with adjacency matrix *A*. For us this is a model of a covalent chemical modification (a “mark”) spreading between monomers of a frozen polymer. Marks spread between adjacent monomers at rate *S* and are lost everywhere at rate *L*.

Let *X_i_* be a random variable which takes the value 1 if the ith monomer is marked and takes the value 0 if the ith monomer is not marked. The expectation value of *X_i_* obeys the differential equation [64]:

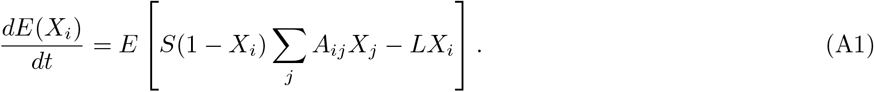

Note that this cannot be turned into an closed equation for *E X_i_* since the *X_i_* are not independent variables in general, *E*(*X_i_X_j_*) ≠ *E*(*X_j_*)*E*(*X_j_*). However, it turns out that it is true (for this model) that *E*(*X_i_X_j_*) ≥ *E X_i_ E*(*X_j_*)—a mark on one monomer can only ever make it *more* likely that another monomer is marked. Therefore,

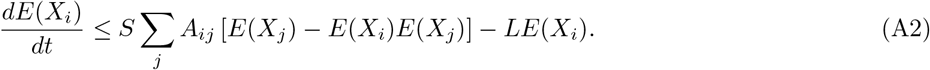

The assumption of equality in the equation above is a mean-field approximation called the *N*-intertwined mean-field approximation (NIMFA) [65, 66]. Importantly, the accuracy of NIMFA appears to increase (e.g. the inequality above becomes closer to equality) as the steady-state mark probabilities *E X_i_* increase [67].

### 1. Regular graph

Now suppose *G* is a regular graph—one where every single node has the same number of neighbors (degree) *d*. Let’s also focus on the *steady-state* of the SIS model, reached after a long time, and suppose that in that steady-state, the probability of any site being marked is the same as that of any other—i.e. *E X_i_* = *p* for all *i*. Then, we have

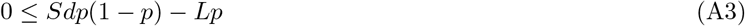

which means that either *p* = 0 or:

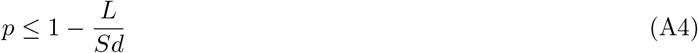

And so according to NIMFA, *p* ⪅ max(1 – *L*/(*Sd*), 0), and this approximation gets better as *ρ* increases.

### 2. A graph with dense and diffuse regions

Now suppose we have a network with a dense region where a typical node has *d*_+_ neighbors and a diffuse region where a typical node has *d*_-_ neighbors.

A very crude approach to understanding the behavior of the steady-state mark probability on this network is to treat these two regions as independent, regular graphs with degrees *d*_+_ and *d*_-_, respectively. Given this (admittedly very strong), simplifying assumption, we would expect that the steady-state mark probability in the dense region satisfies 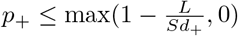 and that in diffuse region we have 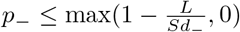.

But, if the dense region is dense *enough*, then NIMFA is a good approximation there, and we have 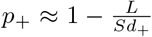 while in the diffuse region we still have 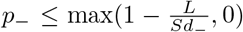. And so, in this regime, there is a range of values of

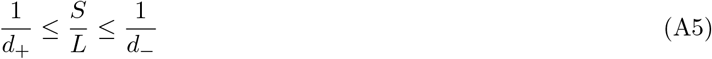

in which there are marks in the dense region but very few (in the approximation, none) in the diffuse region.

## Appendix B: Modeling limited enzyme

Consider a Michaelis-Menten-like enzyme *E* that binds to *AB* pairs and then (once bound) converts them to *BB* pairs, e.g.

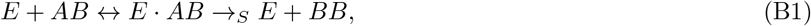

where *E · AB* is an intermediate complex, and *S* is the rate constant for the catalysis step. The instantaneous rate *V_B_* of catalysis (the creation rate of *B*), globally (e.g. in the whole nucleus) is

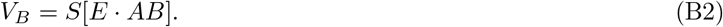

Suppose the formation of the complex is very fast and can be assumed to always be at equilibrium, so that

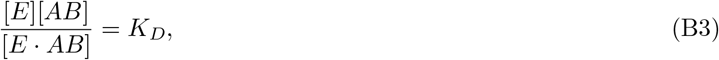

where *K_D_* is a dissociation constant. The total enzyme concentration *E_T_* = [*E*] + [*E · AB*] is always conserved. On the very fast timescale of complex formation, the total number *N_AB_* = [*AB*] + [*E · AB*] of AB pairs is also approximately conserved, which leads to

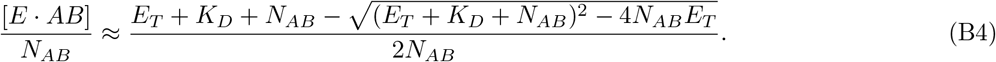

Finally, let’s suppose the binding of the enzyme to AB pairs is very strong, so that *K_D_* → 0. In this limit, we find

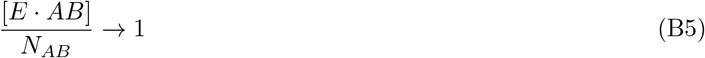

when *N_AB_* ≤ *E_T_*, and

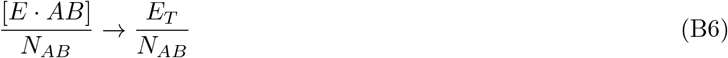

when *N_AB_* > *E_T_*.

Subject to these assumptions of timescale separation and strong enzyme binding, we therefore find that *for any given AB pair*, there is an effective spreading rate *S*_eff_:

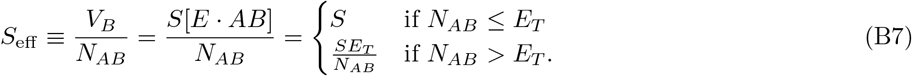

## Appendix C: Supplementary Figures

**FIG. S1.**
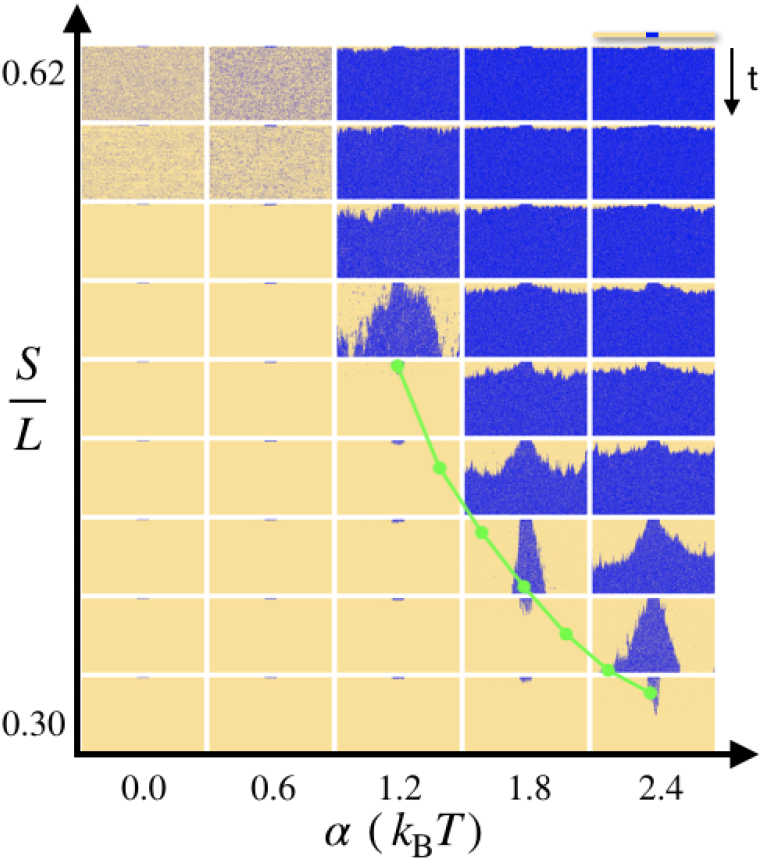
The average value of the ratio *N_B_*/*N_AB_* plotted as a function of *α* (green) in the limited enzyme model, superimposed on Figure 3b, which illustrates the behavior of the unlimited enzyme model for different parameter values. *N_B_*/*N_AB_* appears to lie very close to *λ_c_*(*α*), the critical value of *S/L*, as expected from the argument in the main text.

**FIG. S2.**
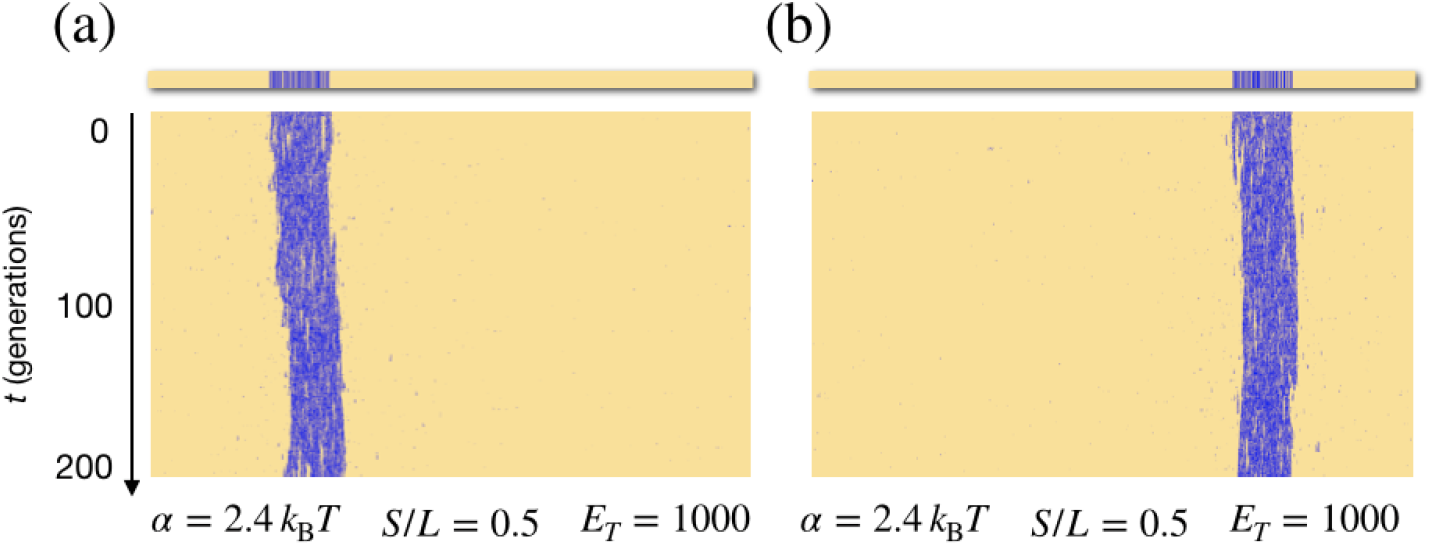
Positional memory. The memory in our model, in the limited enzyme regime, is one of domain *position*. This is illustrated in panels (a) and (b), which show two realizations of our model with identical parameters, but different initial positions of a single contiguous domain of marks. The initial position of the domain is “remembered” for at least hundreds of generations.

**FIG. S3.**
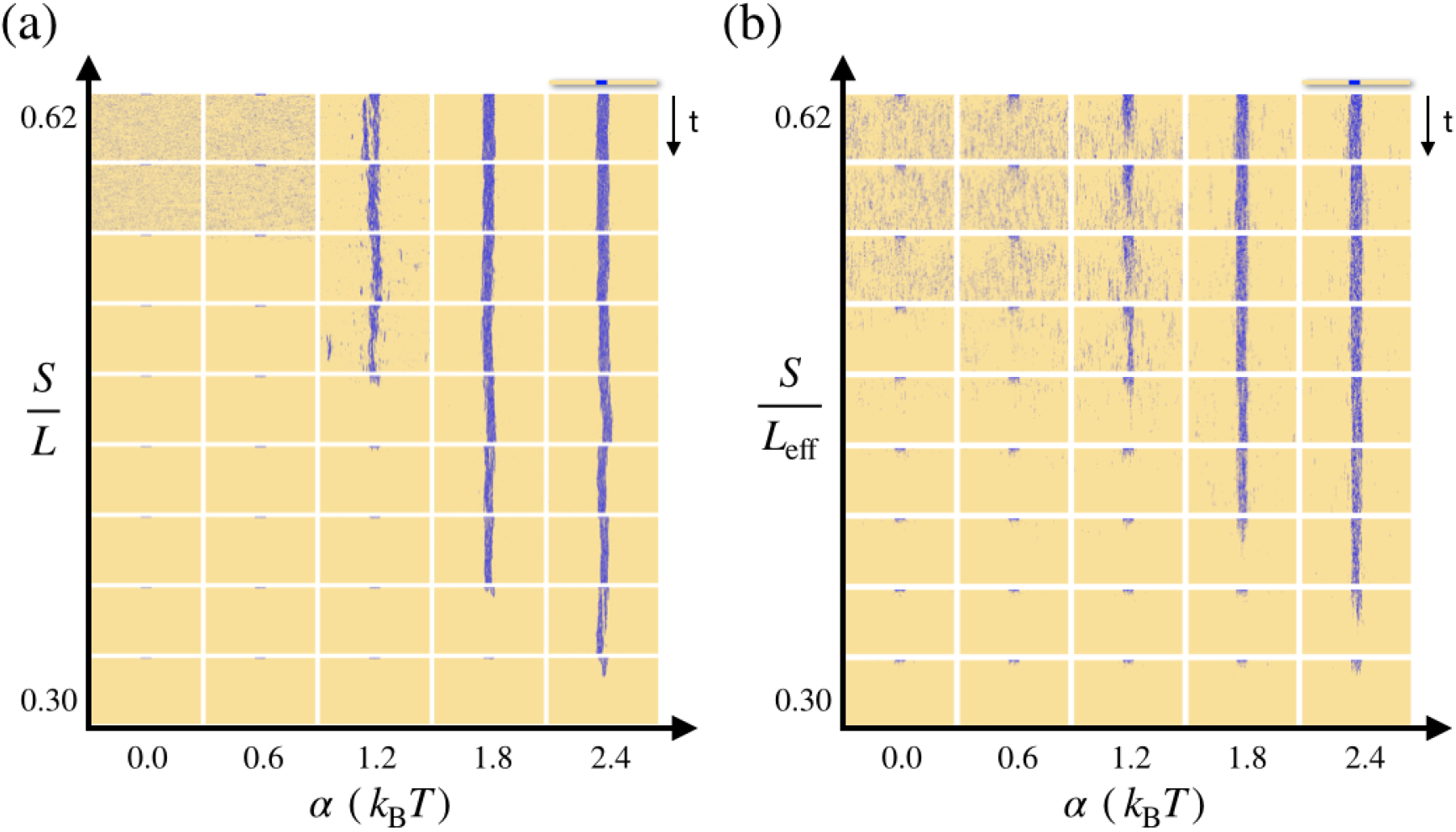
Comparison of the behavior of (a) our model with constant loss of marks at a rate *L* and (b) our model where loss is purely by dilution, periodicially with period *T*_div_. *L*_eff_ = log(2)/*T*_div_.

## Notes

### Competing Interest Statement

The authors have declared no competing interest.

